# A scoping review of the “at-risk” student literature in higher education

**DOI:** 10.1101/2022.07.06.499019

**Authors:** Colin Chibaya, Albert Whata, Kudakwashe Madzima, Godfrey Rudolph, Silas Verkijika, Lucky Makhoere, Moeketsi Mosia

## Abstract

Institutions’ inclination to fulfilling the mandate of producing quality graduates is overwhelming. Insistent petition for institutions to understand their students is about creating equitable opportunities for the diverse student bodies. However, “at-risk” students ubiquitously co-exist. This article conducted a scoping review of literature published locally and internationally that sought to understand “at-risk” students in higher education. The study examined the aims, participants, variables, data analytics tools, and the methods used when the topic on “at-risk” students is studied. Broadly, we sought the bigger picture of what matters, where, when, why, and how so. The Population, Concept, and Context (PCC) framework was considered for demarcating appropriate literature for the *concept* and *context* of “at-risk” students. The JBI protocol was chosen for selecting relevant literature published between 2010 and 2022, searched from the EBSCOhost and ScienceDirect databases. A search tool was developed using the *litsearchr* R package and screening proceeded guided by the PRISMA framework. Although 1961 articles were obtained after applying the search criteria, 84 articles satisfied the stipulated inclusion criteria. Although Africa is lagging, research on “at-risk” students is exponentially growing in America, Europe, and Asia. Notably, relevant articles use academic data to understand students at risk of dropping-out or failing in the first year. Often, statistical and machine learning methods were preferred. Most factors that determined whether a student is at risk of failing or dropping out were found to be highly correlated with high school knowledge. Also, being “at-risk” connoted one’s geographical context, ethnicity, gender, and academic culture. It was noted that autonomously motivated students, with good time management, succeed. Ideally, institutions need to identify areas that need intervention, including courses where special tutoring programmes are needed. Institutions should detect staff who need further training. Nonetheless, psychosocial well-being programmes should augment institutional investments to improve students’ success. Precisely, institutional environments should be stimulating, conducive, and motivating.

## Introduction

Research on “at-risk” students requires one to holistically comprehend this key term, “at-risk” student. Furthermore, it is necessary to conduct an in-depth scoping review of this knowledge domain. This is because the term “at-risk” student should be contextualized and grounded in the factors that distinguish such students from the rest. That understanding may allow institutions to better prepare for new student cohorts in terms of both the necessary resources and infrastructure. At the same time, that understanding may also provide clearer criteria for inclusion and exclusion of the literature that may propel further research in this knowledge domain. Additionally, a broad understanding of the term, “at-risk” student, may potentially provide hints on the appropriate search strategies for related literature, as well as elucidating apt mechanisms for screening suitable studies for the scoping review. In this context, a scoping review is about the synthesis of research that aims to map literature on a topic to the identification of the population of articles, the key concepts, and the context of the knowledge domain thereof [1]. Correspondingly, scoping reviews explicate the evident gaps in the knowledge domain while pinpointing the common characteristics of the evidence thereto, towards informing practice and policymaking.

Our understanding of an “at-risk” student is that of one who would likely dropout, stop-out, burn out, or fail to complete a study programme [3] in higher education. In this context, a dropout is a student who permanently quits from studies without attaining the intended qualification [9]. On the other hand, a stop-out is a student who temporarily discontinues studies with the hope of re-registering at a later stage [9]. Contrary, burning-out is a situation where a student responds to chronic stress through emotional and physical exhaustion characterized by low productivity [14]. Then, failing is a situation where a student endures through a study programme, however, without achieving the desired performance to pass [9]. Understanding the broad literature that characterizes “at-risk” students may inspire focused research for students’ success.

The higher education literature continues to emphasize early intervention as the preeminent way to save “at-risk” students. Evidence is available to support the premise that identifying an “at-risk” student early simplifies the identification of the barriers which the student needs to overcome [2]. In fact, the implementation of individualized support programmes increases the probability of student success, especially when the causal factors for being at risk are correctly identified in time. More so, use of individualized support programmes such as student counselling or peer tutoring allows the sharing of “at-risk” students’ specific risk information which can facilitate proper and timely intervention [39] at a lower cost [40]. As a result, when attempting to understand “at-risk” students, the focus should be on getting to know the student before attempting to solve the underlying challenges. Although direct intervention programmes dominate the list of remedies for being at risk, some literature connotes indirect interventions as tantamount as well, such as the need for the proper sequencing of courses and logical arrangement of the content covered in the courses that put students at risk [41]. Although indirect intervention influences the performance of students, especially in Science, Technology, Engineering, and Mathematics (STEM) degrees [41], proper intervention follows appropriate identification of the main causes of students being at risk. Prevalently, the following categories of indicators are insinuated in higher education; (a) low pre-entry marks [14], poor grade point average [17], muffled interview scores [18], (b) prior experience [17], prior acquaintance with the chosen programme and career goals [19], or prior intention to dropout [21], (c) dwindling performance in tests [17, 38], negative behaviour [21], extended exhaustion levels [19], the general extent of satisfaction with education [21], little effort exerted on tasks [21], poor study skills, as well as poor attendance [11] and participation in class [20]. Other factors that mildly feature when the topic of “at-risk” students is raised include lack of student support strategies [17] at institutional levels, students’ demographics [17], resource allocation [33, 38] at institutional levels, other educational barriers [13], emotional intelligence [34, 35], and learning behaviour [36, 37] of the student. Institutional strategic plans [24, 25, 28] and decision-making approaches [24] are also singled out. It is alluded that institutions that lack proper strategic planning veiledly marginalize [13] and stigmatize [26] “at-risk” students.

Given the propensity to boost student success rates, and the potential benefits of proactive identification of “at-risk” students, most institutions are shifting focus to students’ data for insights. It is our hope that reframing and expanding the concept of an “at-risk” student from data and gaining a better understanding of the underlying scope of work in this knowledge domain would create equitable opportunities for students while also advancing institutional roles in effectively addressing the elements that put students at risk. This scoping review synthesizes research evidence within the “at-risk” student knowledge domain with the goal of mapping the broad concepts to the likely intervention, emphasizing variabilities in the quoted aims, research design strategies, the population of participants, methodological standards, and the reported findings. An especially important point to note is an attempt to fully understand the data upon which the evidence provided is based.

### Objectives

Three objectives summarize this scoping review in the sequence they are presented as follows: (a) We want to identify articles that present prevalent categories of “at-risk” students in the higher education context. (b) We also want to investigate the prevalent aims, data analytics tools, common participants, variables, and methods insinuated when the topic of “at-risk” students is being studied. Last, (c) we want to analyze the articles that meet the inclusion criteria to obtain a broader picture of what matters, where, when, why, and how the problem of the “at-risk” student has been tackled in the past. Achievement of these objectives may give insights to guide further studies aimed at bringing about change and social justice in higher education.

### Research questions

Three questions are asked in line with the objectives as follows; (a) Which articles tackled the “at-risk” student problem in the higher education context? (b) What were the aims, data analytics tools, participants, variables, and methods used in tackling the problem? (c) What mattered, where, when, why, and how was the “at-risk” student problem addressed? Hopefully, answers to these questions may provide intuition into further research to guide practices and propel data-driven institutional planning.

### Overview

The rest of the article proceeded as follows; a section on how the PCC framework fits into this study follows next. The PCC framework guides the selection of the *population* of articles that befit the *concept* and *context* of the study. The methods we followed in completing the study are presented thereafter, emphasizing the inclusion and exclusion criteria, search strategy, screening procedure, and how the summaries were drawn. Subsequently, the results which report the distribution of articles followed before the conclusion highlighted the contributions and direction for further studies.

## The PCC framework

This scoping review categorized articles on the “at-risk” student in higher education. An appropriate search strategy for articles published on this topic was proposed. In this case, we adopted an a priori model known as the PCC (*Population, Concept*, and *Context*) framework [31], asking the following question: *“Which literature seeks to understand the “at-risk”student knowledge domain in the higher education context?”*. This PCC framework shows a plan for what matters [31] in an open *population* of articles. It would imply that all articles that mention the *concept* of an “at-risk” student may be included. However, the inclusion criteria define the boundary of articles that fit into the desired *population, concept*, and *context* of the study. Precisely, the key *concept* remained the “at-risk” student. This is a broad concept that could cover any kind of articles that mention the term, “at-risk” student. However, the PCC framework was used to contextualize the *concept* of “at-risk” students through a clearly defined search strategy that stipulated how the relevant articles were selected and screened, bearing in mind the higher education setting. Also, the *concept* of an “at-risk” student has been left open regarding the sources of evidence, which may come from anywhere, including the articles where students may be at risk of dropping-out, stopping-out, burning-out, or failing. This scoping review demarcated the *concept* of an “at-risk” student to comprise dropouts, stop-outs, burn outs, and failing students in the higher education perspective. The methods section will meticulously elucidate the *population* and the type of evidence considered in characterizing the *concept* and *context* of this study. Anticipated results were reported using figures and charts that depict the distribution of articles categorized by year, region, aim, participants, methods, data analytics tools, and findings.

## Methods

The inclusion and exclusion criteria, search strategy, method for screening articles, and the ways in which results are summarized are the main sub-sections of this section. The ethical considerations undertaken before the start of this study are also discussed to justify the integrity of the work.

### Inclusion and exclusion criteria

The Joanna Briggs Institute (JBI) scoping review protocol [8, 31] was adopted, where articles with the keywords such as intervention, at-risk, failing, dropout, stop-out, burn out, performance, and success were nominated to define the *population* of articles for this scoping review. The key *concept* remained the term “at-risk” student. Conversely, the *context* was persistently about the characterization of students at risk of dropping out, stopping out, burning out, or failing in higher education. In this case, higher education refers to university and college education. The literature looked at academic performance as one of the most important factors in determining success [27]. The *population* of the articles comprised peer-reviewed conference and journal articles. Only articles that were published in English were considered in this scoping review. A literature search was conducted in the EBSCOhost and ScienceDirect online databases, soliciting articles published between 2010 and 2022.. The deep inner type of the study was not of interest. Therefore, review articles, conceptual papers, theoretical articles, as well as empirical quantitative and qualitative studies all qualified. An iterative approach which allowed repeated refinement of the inclusion and exclusion criteria was adopted. Thus, articles went through several iterated screening rounds before the final list of relevant literature was generated. Disputed articles were considered through consensus after round robin reviews by the research team members. Sometime, detailed manual scrutiny of the full texts of the articles were considered as the last resort.

### Search strategy

A three-step search strategy was employed. First, we employed the *litsearchr* R package [28] as a tool to facilitate a quick, objective, and reproducible search using text-mining and keyword co-occurrence networks <u>[28].</u> This approach reduced possible bias in the search by removing the reliance on predetermined factors. The tool improved search recall by exploiting the identification of synonymous terms that research team members would otherwise miss. Also, it took away the likely bias of researchers typically selecting keywords based on their own knowledge without specifying how the search process was administered [10]. Such bias would instigate irreproducibility because it would be hard to recall the procedure followed in each selection of a comprehensive set of concepts. The following search query was used to mine the relevant articles.

*(students AND at-risk AND (failing OR stop out OR burn out OR dropout) AND (university OR college))*

The validity of this search query was verified with the help of an experienced librarian. Consultations with content experts in the field of student success were also considered to triangulate the search strategy, as well as to enhance rigour and reliability. In this case, content experts were a valuable resource for finding literature that was hard to identify through other means. The second step was about the actual search process, where the search query was executed following the directions from content experts. The final step focused on scrutinizing the list that passed the inclusion criteria for any outstanding patterns.

### Screening of included articles

The standard procedure to verify scientific material is through manual screening. Generally, such screening can be split into several steps, including screening articles by titles, screening by abstracts, or screening by physically going through the full text. The *revtools* [16] R package that supports evidence synthesis was considered for the first round of screening. This tool de-duplicates bibliographic data using titles and abstracts. It also visualizes articles using topic models, allowing articles to be screened by removing duplications arising from using different search strategies. In using the *revtools* R package, we were guided by the PRISMA-ScR framework [15]. PRISMA stands for <u>P</u>referred <u>R</u>eporting Items for <u>S</u>ystematic reviews and <u>M</u>eta-<u>A</u>nalyses [15] while ScR is an acronym for Scoping Review. The PRISMA framework facilitated the construction of a flow diagram that shows how screening was undertaken through the different stages of the scoping review, reporting the articles considered and those excluded, together with the reasons for the exclusion.

### Summaries

A standardized data extraction template that followed the PRISMA-ScR format was created as part of the data charting process. We indicated that the *population* of articles that met the inclusion criteria for the *concept* of the “at-risk” student in the *context* of failing, dropping-out, stopping-out, or burning-out in higher education, together with the details of those articles in terms of the year of publication, country, aim, participants, methodology, intervention, and findings, were the key results reported and analyzed in this scoping review. We mainly looked at the characteristics of these articles to establish likely knowledge gaps to explore further. We also sought the bigger picture of what matters, where, when, why, and how literature tackled the “at-risk” student problem. Figures, charts, and tables were the main reporting tools [12] used because they better depict the gap maps in the knowledge domain under study.

### Ethical statement

The research protocol for this study underwent approval by the Senate Research Ethics Committee of the Sol Plaatje University. The work was endorsed by the Directorate of the Centre for Teaching, Learning, and Programme Development. The larger project, from which this study ensued, is registered at the National Teaching Advancement Programme as an institutional project. Hopefully, the results from the project will instigate change and social justice in higher education and inform further research on good practices towards data-driven institutional planning and decision-making.

### Search Results

Figure 1 shows the PRISMA-ScR flow diagram that summarizes the articles considered, included, and excluded. The PRISMA-ScR seeks to determine the articles that tackled the “at-risk” student problem in the higher education context (research question (a)). Precisely, 1918 articles that were extracted from the ScienceDirect and EBSCOhost databases using the proposed search query made it through the first round of inclusion. An additional 43 articles qualified through random search from the internet (28 articles), or from recommendations by content experts (3 articles), and citation search (12 articles). A total of 1961 articles, thus, formed the desired scoping review *population*. The initial screening process using the titles of the articles dropped 139 articles because they had duplicate titles. Further screening using the abstracts excluded another five articles. Six more articles were discarded because their topics were not in line with the scope of the key *concept* and the *context* of the study. Therefore, the *population* of relevant articles dropped to 1768. The application of the *revtools* [16] R package eliminated the largest chunk (1560 articles) through topic modelling, title, and abstract screening. The remaining 220 articles were reviewed manually. However, the full texts for 53 of the 220 articles could not be retrieved, thus reducing the number of articles to 167 articles. These 167 articles were subjected to additional manual screening to check whether their content was in line with the concept of “at risk” students. Another 27 articles were discarded as their participants were not part of the higher education domain. Eleven articles were removed because they focused on the *context* of nursing students in nondegree-offering colleges. There are 6 non-English articles that were also removed. A further 13 articles were dropped because they focused on other *contexts*, such as the risk of quitting or stopping medication or some other programs not related to education. The full-text reviews excluded another seven articles that were identified as duplicates that were missed by the *revtools* automated application tool.ls.

**Figure 1:**
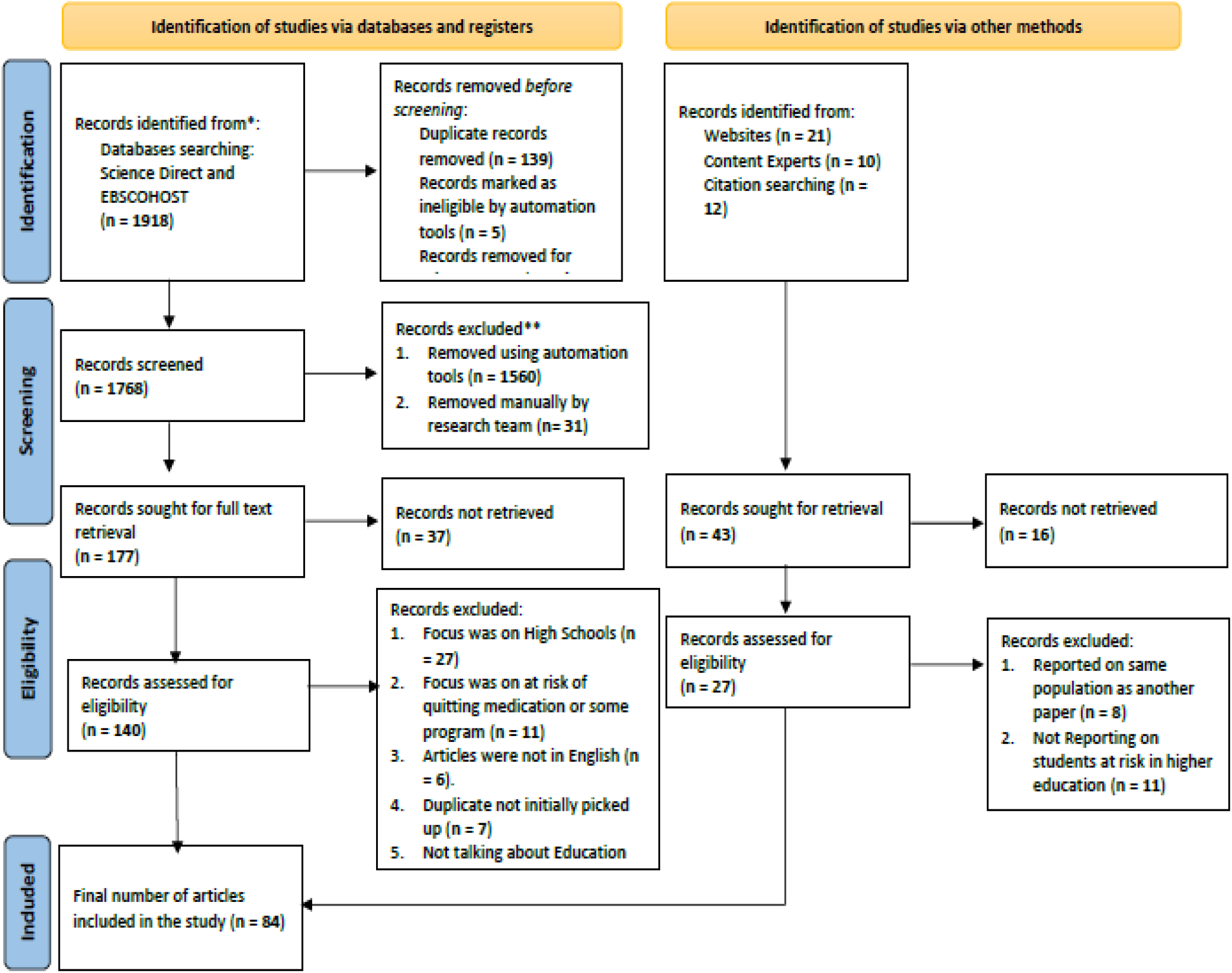
PRISMA-ScR flow diagram.

Additionally, eight articles were eliminated because they reported on the same participants as reported in other articles under consideration. Finally, 11 articles were removed because they differed in their definition of “at-risk” students. Eventually, 84 articles remained as reflected in Figure 1. These articles formed the basis of the findings, discussions, recommendations, and conclusions that are reported in this study.

### Findings, Discussions, and Recommendations

Literature notes the prevalent factors/variables that determine whether a student is at-risk include marks [17, 18], prior learning experience and the student’s prior intentions [17, 19, 21], pre-entry expectations [30], student’s personal behaviour [11, 17, 20, 21, 38], and partly the social environment of the student [17]. Purportedly, the presence of these factors/variables likely suggests poor outcomes in the student’s future [11]. It is also insinuated that an “at-risk” student would predominantly demonstrate challenges with internalization and externalization of learning content [11]. Even compelling is the observation that an “at-risk” student would surely require intervention programs for success [19]. Such interventions should dominantly revolve around peer mentorship, tutorship, group studies, and enhanced residence culture. Little is said regarding non-academic factors such as students’ socio-economic factors, childhood experiences, or family careers [29].

Figure 2 is a snapshot of the popular terms used to characterize “at-risk” students, with the terms such as dropout, poor performance, at-risk, and success standing out. This observation is in line with the views that ensue from topic modelling of the dominant variables used to identify the top “trending topics” on the “at-risk” student.

**Figure 2:**
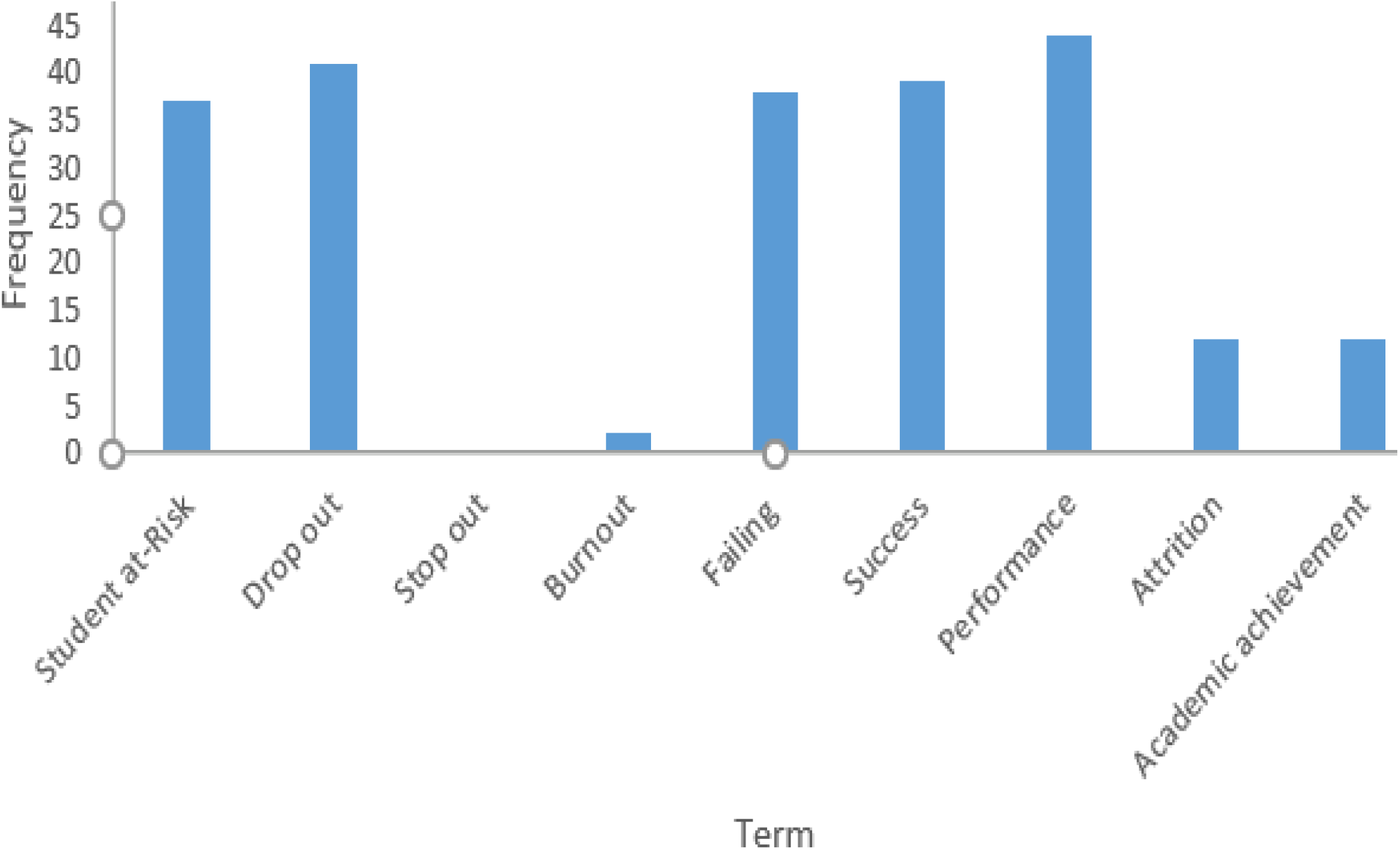
Popularity of terms used in the context of “at-risk” students.

The research findings reported in this section sought to determine the aims, data analytics tools, participants, variables, and methods that were employed in the “at-risk” student’s literature. The aim(s) of most articles was to determine the factors/variables that cause a student to be at risk. Table 1 categorizes these top trending topics and factors/variables in the “at-risk” student’s literature. The summaries show that the included articles varied widely in terms of the terminology used to describe these dominant variables. For example, the category “Grades” included factors such as final exam grades, exam scores, major test marks, marks in formative tests, predicted grades, and prior grades. The category “Academic” included factors like academic record, academic motivation, academic support, academic success, academic performance, academic background, and academic integration. Literature indicates higher occurrences of the terms: Grades (53.8%), Academic (28.6%), Gender (18.7%), GPA (13.2%), Age (12.1%), Data (11%), Course (9.9%), Race/ethnicity (9.9%), Study (7.7%), Support (7.7%), Time (7.7%), Semester (6.6%), Scores (5.5%), Education (5.5%), and Parent (5.5%). This observation is consistent with the marks being a common factor of “at-risk” students [17, 18].

**Table 1:** Top trending categories of the variables.

| Category | (N = 84) | Factors/Variables |
| --- | --- | --- |
| **Grades** | 53.8 | Final exam grades, exam scores, major test marks, marks in formative tests, predicted grades, prior grades, grades in core courses, expected course grade, grade(s), secondary school grades, quiz scores, exam scores, homework scores |
| **Academic** | 28.6 | Academic records, academic motivation, academic support, academic success, academic performance, academic background, academic integration |
| **Gender** | 18.7 | Sex |
| **GPA** | 13.2 | GPA |
| **Age** | 12.1 | Age |
| **Data** | 11.0 | Learner and learning data, administrative data, enrolment data, activity data, trace data, records' system data, learning management data |
| **Course** | 9.9 | Course code, course load, key courses, course observations, course non-completion, course credits, course status, Expected course grade, professor of the course, core courses |
| **Race/ethnicity** | 9.9 | Race, race/ethnicity |
| **Study** | 7.7 | Study time, study skills, study program, field of study, study results, study group, work-study |
| **Support** | 7.7 | Educational support, peer support, parental support, family educational support, extra educational support |
| **Time** | 7.7 | Interaction time with content, free time, time management, study time, travel time |
| **Semester** | 6.6 | End of semester survey, semester enrolled, |
| **Scores** | 5.5 | SAT scores, ACT scores, University Entry scores, |
| **Education** | 5.5 | Prior education, education values, education system, prior schooling |
| **Parent** | 5.5 | Parent relationships, parent occupation, parent education |

On the other hand, Figure 3 depicts the distribution of the articles included in this scoping review that were published between the years 2010 to 2022. Generally, efforts to understand the “at-risk” student is exponentially growing. This growth may be attributed to the institutional goodwill earned from understanding students. Institutions that seek to understand their students often achieve good student success rates [17]. They plan better and make data-informed decisions. However, much attention to this bona fide agenda is, notably, visible around Europe, America, and Asia. Regions such as Africa are lagging (see Figure 4a), ostensibly calling for immediate attention.

**Figure 3:**
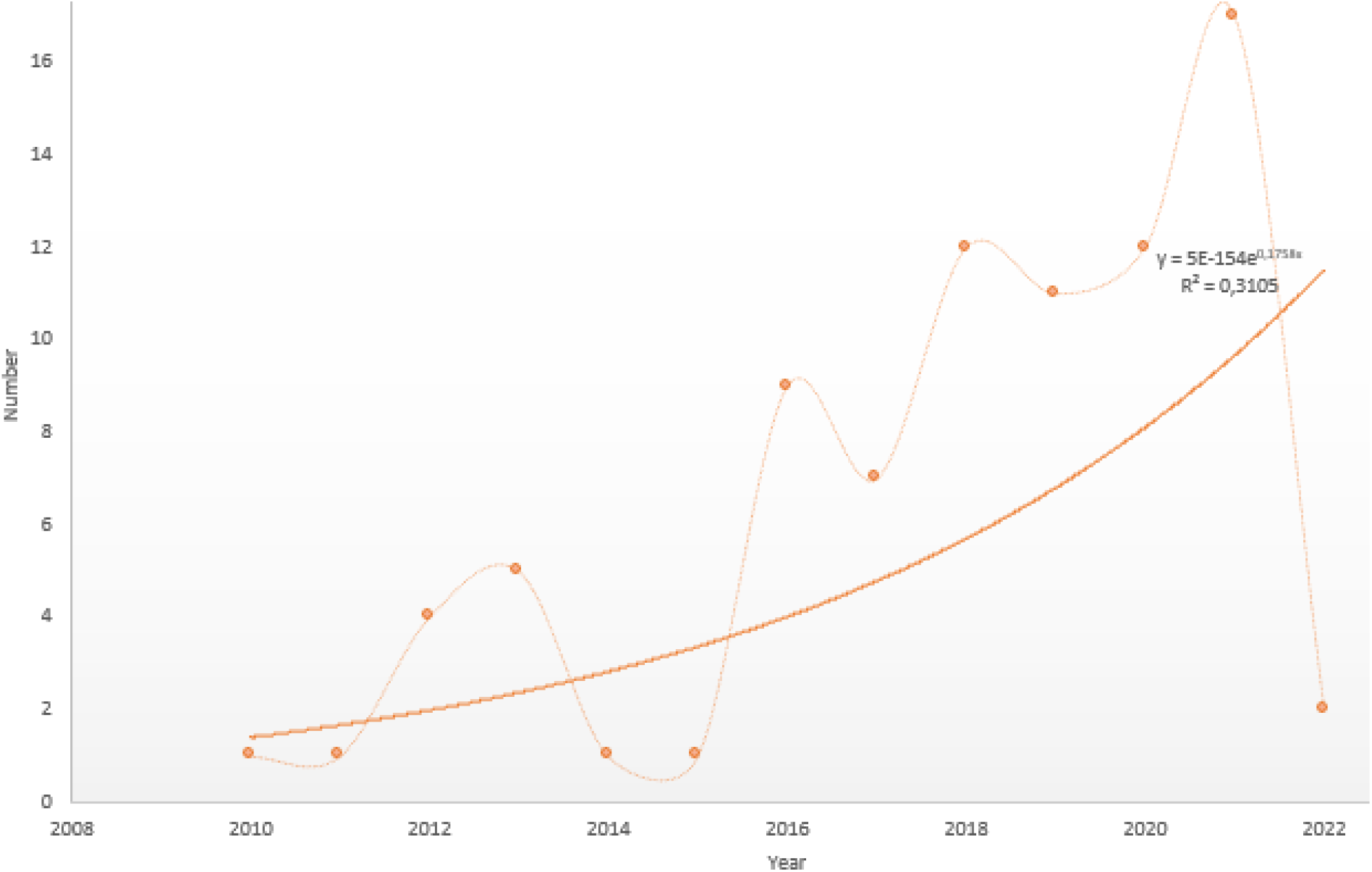
Articles on the “at-risk” students that were published per year between the years 2010 and 2022.

**Figure 4a:**
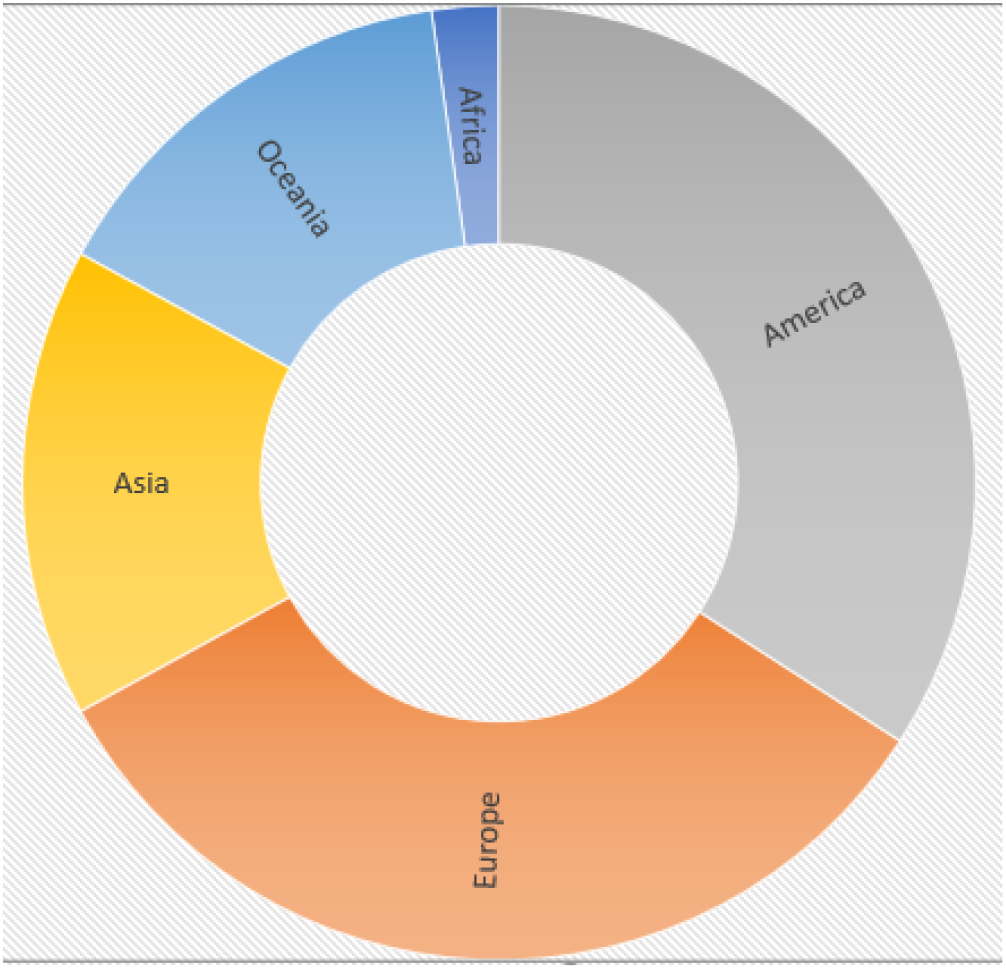
Distribution of articles by continent.

**Figure 4b:**
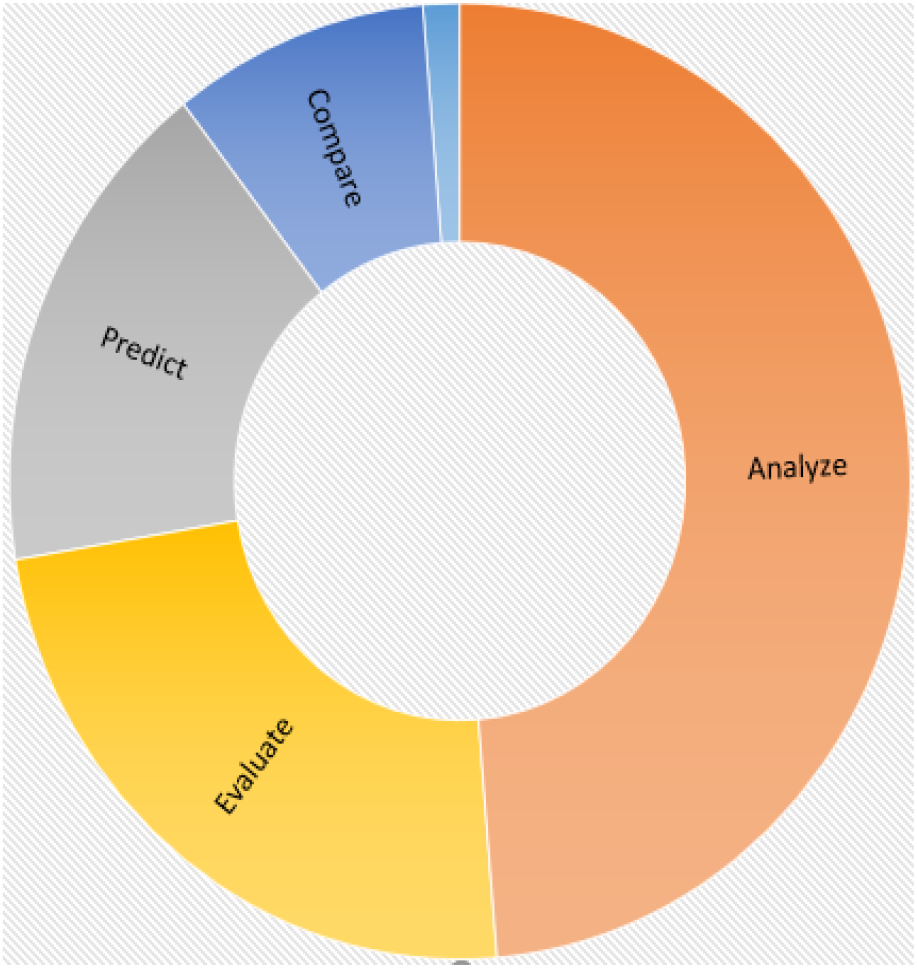
Distribution of articles by aim.

**Figure 4c:**
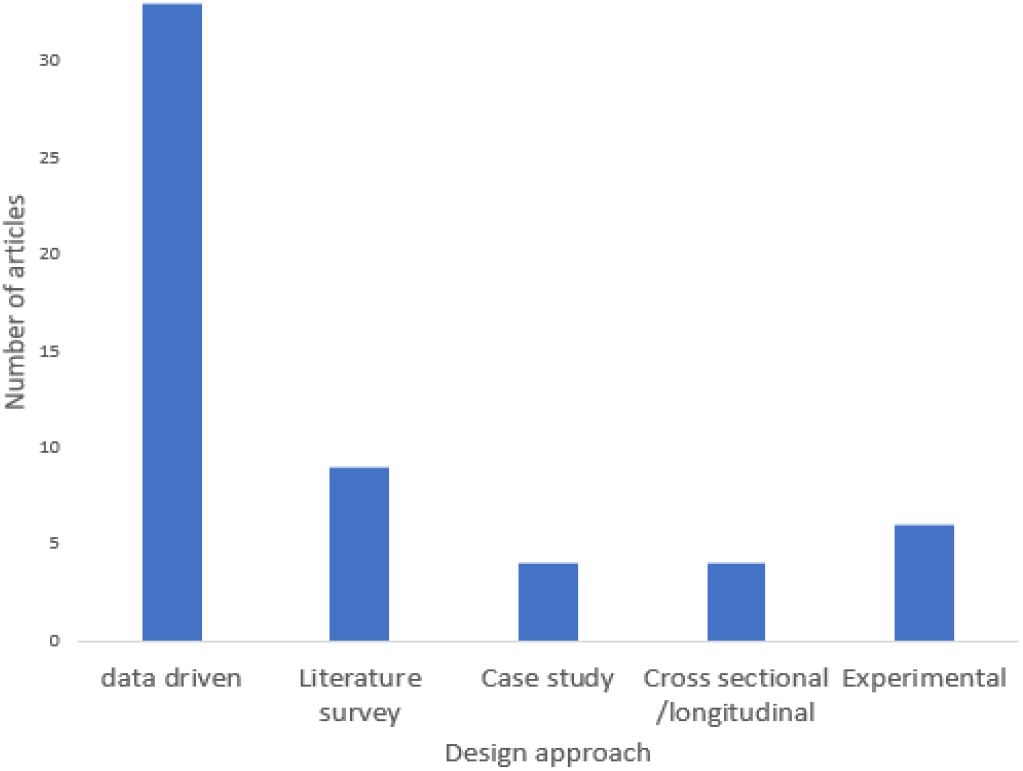
Distribution of articles by study design.

**Figure 4d:**
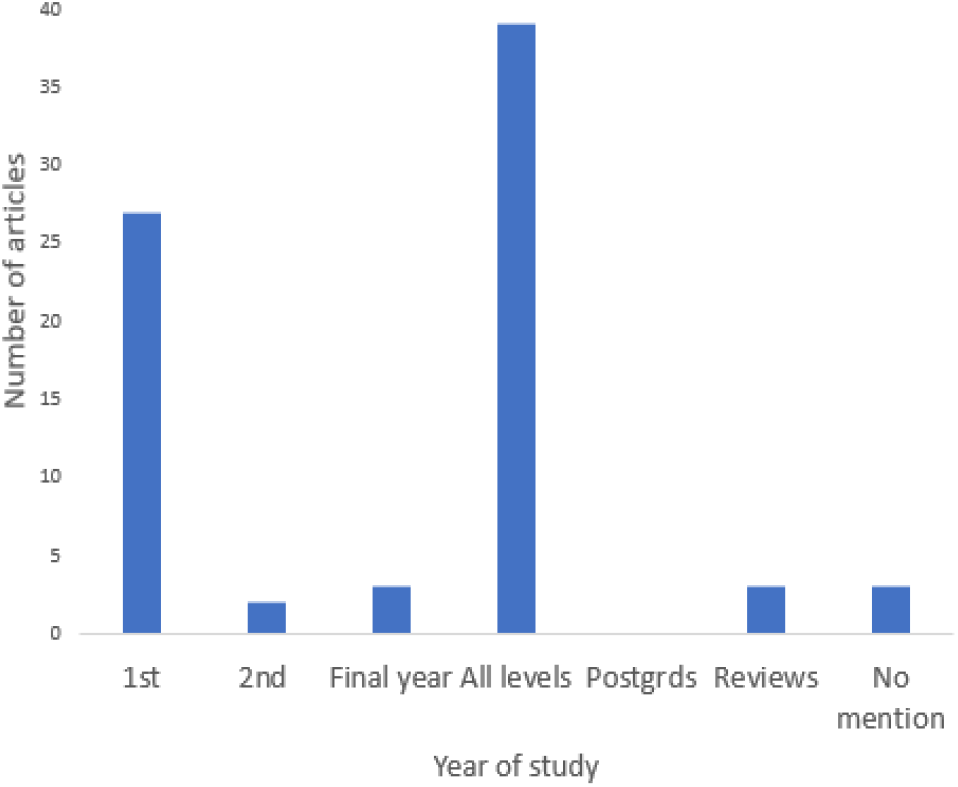
Distribution of articles by participant level.

Most articles included in this study emphasized data analysis to seek institutional advancement towards students’ retention and success at undergraduate level. There is barely any literature on the “at-risk” student in post-graduate studies and that alone is a gap to explore further. First year students are the common target group of participants unless all students in the *context* were considered (see Figure 4d). This may be because cohorts of first year students often comprise the highest number of “at-risk” students. Another reason may be that the transition from high school to university is commonly perceived as radical, which renders first year students as indigent for support than senior students.

Equally, although a good chunk of literature focused on the building of predictive models to identify “at-risk” students, comparative studies to evaluate which model gives plausible outcomes are few (see Figure 4b). This may be because this knowledge domain is still in its infancy and such comparative studies may be upcoming. Nevertheless, data-driven methods are still preferred because of the insights drawn from several data-analytics tools. Studies that focused on surveys, case studies, experimental and cross-sectional research are also quite visible in the literature (see Figure 4c). However, advanced data-analytics models are preferred for simplifying the way meanings can be drawn from the data collected from the many information systems institutions often subscribe to, and this will remain the likely trend in most related future studies. Such data analytics tools are summarized into four broad categories as shown in Figure 5.

**Figure 5:**
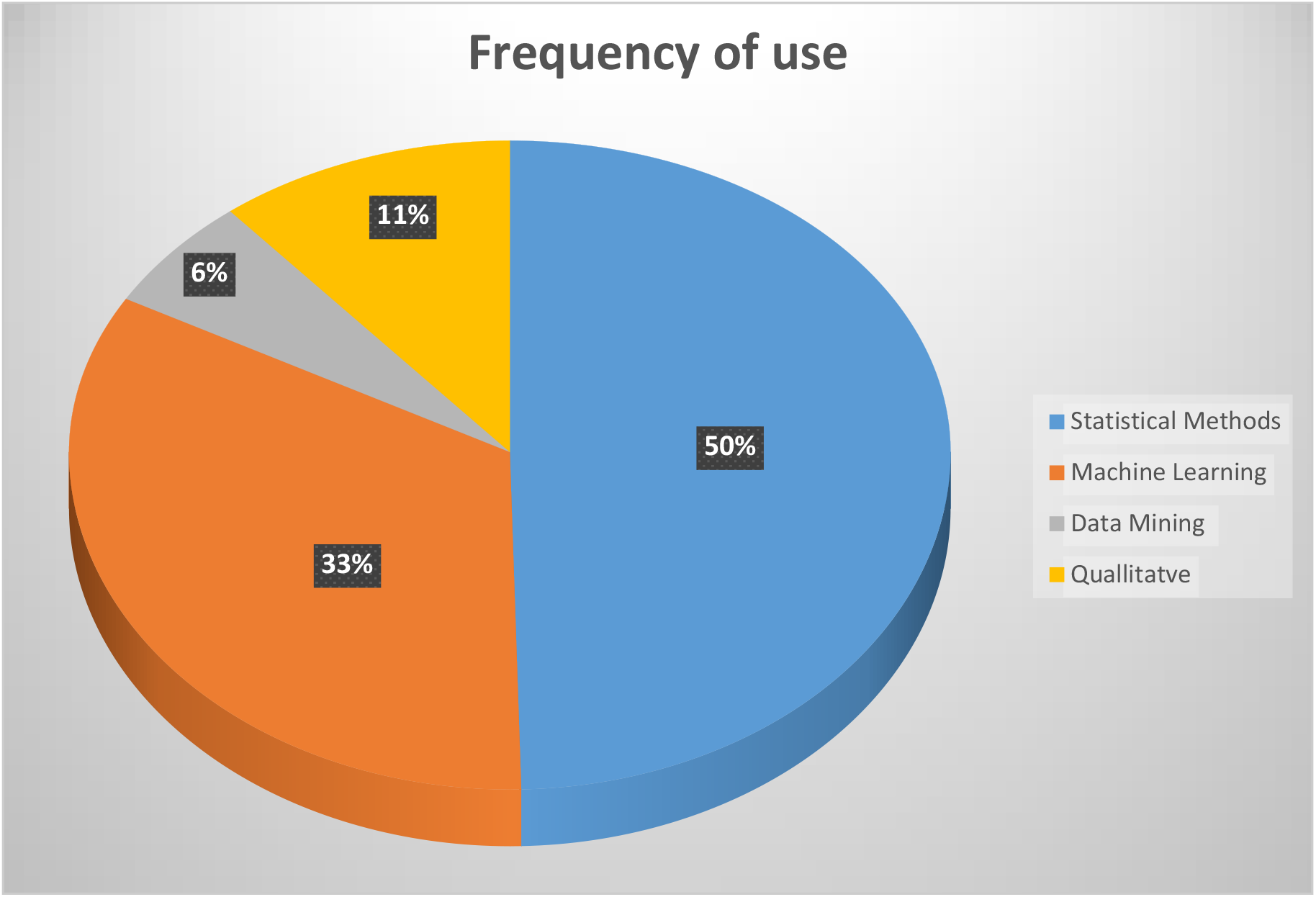
Distribution of methods/data analytical tools used in the articles.

Some articles employed more than one data analytics tool. Statistical methods were preferred most and were used for approximately 50 % of the time. Examples of statistical methods included survival analysis, confirmatory analysis, descriptive statistics, logistic regression, multiple linear regression, cox regression, and analysis of variance. On the other hand, machine learning methods are also quite dominant, employed for approximately 33% of the time. Some of the examples of the machine learning methods preferred include decision trees, artificial neural networks, naive Bayes, K-nearest neighbour, support-vector-machines (SVM), and different Ensemble methods. In addition, data mining techniques and qualitative methods also feature quite frequently (used for about 6 % and 12 % respectively).

Several findings emanated from the scoping review regarding “at-risk” students. Generally, it is repeatedly insinuated that students will likely dropout if their secondary school knowledge was low or their motivation to study was low [42]. That intrinsic engagement reduces the chance of the burn out syndrome. Positive personality and commitment, coupled with determinants of cognitive skills, attest the impact of that conscientiousness against dropping out. Autonomous motivation and good time management are positive predictors of achievement. High correlations are alluded between high school knowledge and dropout intention, satisfaction with education, academic exhaustion, and the student’s expectations of graduation [4]. Low dropout rates were also linked to students who participated in social groups [43]. However, funding challenges then implicated the influence of geographical location and ethnicity as indicators of “at-risk” students.

Intervention close to individualized attention are seen as more effective, including peer tutoring and one-on-one counselling. Subscription to the use of early warning systems that reduce the burden of counseling, systems that will work towards enhancing metacognitive awareness, self-awareness, and self-regulation, as well as tracing logs by students on learning management systems may also simplify early prediction of “at-risk” students. Most compelling is the need for institutions to identify courses that are hard-to-pass and evaluate the question papers to determine the levels of difficulty. Lecturers should also implement student motivation strategies, including provision of timely feedback on assignments. Interventions that focus on the psychosocial well-being of students and the emotional intelligence of students are also recommended. Machine learning models such as AutoML can be adopted to formulate optimal student performance prediction models that use pre-start data. More interpretable models that provide educators with course feedback on student status are also recommended. Creation of caring, supportive, and welcoming environments within the university is critical to creating that sense of belonging.

### Gaps to explore

The topic of “at-risk” student is receiving close attention. However, focus to the different arms of the *concept* of an “at-risk” student is not fairly spread. Emphasis is tilted towards interventions against dropping out or failing. Little is visible regarding students at risk of stopping-out or burning out and that is an apparent avenue for further studies in this body of knowledge. Similarly, most articles dwelt on the *concept* of an “at-risk” student in the *context* of dropping out or failing from American, European, or Asian institutions. Studies on this *concept* in African institutions’ perspectives are rare. Research to compare the results yielded with the context of African institutions is worthwhile. Such studies may take us closer to the generalized understanding of an “at-risk” student beyond undergraduate levels. More so, literature suggests that students’ internal states are also predictors of performance. Data about student’s prior experiences, social interactions, relationships, and extracurricular activities is, thus, needed to further inform the understanding sought. A gap spins around investigating the use of non-academic data to define students’ journeys [44]. Lastly, little is also said about the evaluation of the proposed interventions. Not much is known about the effectiveness of the interventions and that alone, is also a gap worth undertaking.

## Conclusion

Unfolding the population of articles that characterize “at-risk” students guided the aims, participants, methods, variables, interventions, and data analytics tools one can adopt in related studies. Three contributions apparently stand out as follows:

- The scoping review set forth an understanding of the *population of studies, concepts*, and *context* of the “at-risk” student.
- Institutions of higher learning can build on this understanding to similarly get to know their own diverse student bodies.
- The scoping review elucidated various applications of different data analytics tools in understanding “at-risk” students.
- Tailored studies which suit particular scenarios may ensue.
- Although the focus of this scoping review was on understanding the “at-risk” student in the higher education space, the results presented create a baseline context upon which a broader understanding of students, in general, may emanate.

A few challenges are observed from this scoping review as follows:

- Although scoping reviews comprehensively synthesize evidence, dealing with a broad range of literature may blur important methodological steps which makes it difficult to establish boundaries.
- A good scoping process requires more time and resources that are often difficult to predict at the start of the research.
- Crafting an appropriately inclusive search query which would drop the number of screening iteration is hard.
- Manually assessing the validity of some of the articles to be included when disputes arise is even harder.

Four ambitious directions for future work are envisioned as follows:

- Investigations to corroborate the “at-risk” student knowledge domain to the African context are apparently overdue.
- This scoping review could be enriched by extending the *context* of the study to accommodate other use cases.
- Further research is paramount which analyzes trace data to better understand the broader spectrum of the enrolled student
- It is worth checking the extensibility of the *concept* of “at-risk” students to include demographic and institutional aspects

## Acknowledgments

This work was undertaken under the leadership of the DVC’s office, guided by the Centre for Teaching, Learning, and Programme Development of the Sol Plaatje University, supported by the Department of Higher Education and Training, through its University Capacity Development grant. We acknowledge their financial and moral support.

